# Genetic background modifies vulnerability to glaucoma related phenotypes in *Lmx1b* mutant mice

**DOI:** 10.1101/2020.07.05.188516

**Authors:** NG Tolman, DG Macalinao, AL Kearney, KH MacNicoll, CL Montgomery, WN de Vries, IJ Jackson, SH Cross, K Kizhatil, KS Nair, SWM John

## Abstract

Variants in the LIM homeobox transcription factor 1-beta gene (*LMX1B*) predispose individuals to elevated intraocular pressure (IOP), a key risk factor for glaucoma. However, the effect of *LMX1B* mutations varies widely between individuals. To better understand mechanisms underlying LMX1B-related phenotypes and individual differences, we backcrossed the *Lmx1b*^*V265D*^ (also known as *Lmx1b*^*Icst*^) allele onto the C57BL/6J (B6), 129/Sj (129), C3A/BLiA-*Pde6b*^*+*^/J (C3H), and DBA/2J-*Gpnmb*^*+*^ (D2-G) strain backgrounds. Strain background had a significant effect on the onset and severity of ocular phenotypes in *Lmx1b*^*V265D/+*^ mutant mice. Mice of the B6 background were the most susceptible to developing elevated IOP, severe anterior segment developmental anomalies (including malformed eccentric pupils, iridocorneal strands, and corneal abnormalities) and glaucomatous nerve damage. In contrast, *Lmx1b*^*V265D*^ mice of the 129 background were the most resistant to developing anterior segment abnormalities, had less severe IOP elevation than B6 mutants at young ages, and showed no detectable nerve damage. To identify genetic modifiers of susceptibility to *Lmx1b*^*V265D*^-induced glaucoma-associated phenotypes, we performed a mapping cross between mice of the B6 (susceptible) and 129 (resistant) backgrounds. We identified a modifier locus on Chromosome 18, with the 129 allele(s) substantially lessening severity of ocular phenotypes, as confirmed by congenic analysis. By demonstrating a clear effect of genetic background in modulating *Lmx1b*-induced phenotypes, by providing a panel of strains with different phenotypic severities and by identifying a modifier locus, this study lays a foundation for better understanding the roles of LMX1B in glaucoma with the goal of developing new treatments.

## Introduction

Glaucoma is a group of complex disorders that share a characteristic pattern of visual field deficits and retinal ganglion cell degeneration. It is a leading cause of blindness worldwide affecting 80 million people (Quigley and Broman, 2006). Important risk factors for glaucoma include elevated intraocular pressure (IOP), genetics, and advanced age. Lowering IOP to a safe level is the only available treatment (Weinreb et al., 2014). The aqueous humor (AqH) drainage tissues, including the Schlemm’s canal (SC) and trabecular meshwork (TM), have a key role in controlling IOP (Fautsch and Johnson, 2006). Resistance to AqH drainage from the eye through the SC and TM is important in determining IOP. However, the mechanisms underlying dysfunctional AqH drainage and subsequent IOP elevation require additional characterization. A majority of glaucoma cases are attributed to primary open angle glaucoma (POAG), where IOP elevation lacks an obvious physical cause (Quigley and Broman, 2006). Recently, genome wide association studies (GWAS) have improved understanding of the genetic basis of POAG by implicating more than 70 loci (Bonnemaijer et al., 2018; Choquet et al., 2018; Choquet et al., 2020; Genetics of Glaucoma in People of African Descent et al., 2019; Khawaja et al., 2018; MacGregor et al., 2018; Taylor et al., 2019; Youngblood et al., 2019). Research that defines how these genes affect IOP is expected to yield new drug targets and improved treatments for lowering IOP (Choquet et al., 2020).

The LIM homeobox transcription factor 1-beta (*LMX1B*) gene was associated with elevated IOP and POAG through genome wide association studies (GWAS) and has been validated in multiple populations (Choquet et al., 2018; Gao et al., 2018; Gharahkhani et al., 2018; Khawaja et al., 2018; MacGregor et al., 2018; Shiga et al., 2018). Prior to GWAS, dominant mutations in *LMX1B* were identified to cause nail-patella syndrome (NPS) (Chen et al., 1998; Dreyer et al., 1998; Vollrath et al., 1998). NPS is a developmental disorder with characteristic symptoms including nail dysplasia and abnormally developed limb structures (Farley et al., 1999; Sweeney et al., 2003). Within NPS patients, 20-30% develop elevated IOP and POAG, a prevalence that is significantly higher than in the general population (Mimiwati et al., 2006; Sweeney et al., 2003). Apart from POAG, there are a wide range of ocular phenotypes reported in NPS patients including developmental iris, corneal, and pupillary abnormalities and congenital glaucoma (Lichter et al., 1997; Sawamura et al., 2014; Spitalny and Fenske, 1970). Importantly, there are striking differences in onset and severity of phenotypes between patients that inherit the same *LMX1B* variant (Knoers et al., 2000; McIntosh et al., 2005; Sweeney et al., 2003). Thus, it is likely that genetic background modulates the risk of developing specific disease phenotypes in patients with *LMX1B* variants. Identifying genetic modifiers of *LMX1B*-related phenotypes will be important in understanding risk of glaucoma and is expected to provide novel mechanistic information on the etiology of IOP elevation.

Based on high sequence homology, mice have been used to understand the biological role of LMX1B in several tissues (McIntosh et al., 2005). Previous work in mice has shown that LMX1B is required for the development and function of AqH drainage tissue including the TM (Liu and Johnson, 2010; Pressman et al., 2000). Mice with dominant point mutations in *Lmx1b* recapitulate several phenotypes found in humans with *LMX1B* variants. One important mutation causes a valine to aspartic acid substitution (*Lmx1b*^*V265D*^, also known as *Lmx1b*^*Icst*^) in the transcription factor’s homeodomain, disrupting its ability to bind DNA (Cross et al., 2014). Mice heterozygous for *Lmx1b*^*V265D/+*^ develop elevated IOP and glaucomatous neurodegeneration (Cross et al., 2014). These mice also present with several additional ocular phenotypes including abnormal SC and TM, congenital defects of the iris such as iridocorneal strands, abnormally open pupils, and corneal phenotypes including corneal opacities, corneal neovascularization, and corneal scarring (Cross et al., 2014). Congenital abnormalities of the iris, cornea, and pupil have also been observed in a subset of NPS patients (Beals and Eckhardt, 1969; Bennett et al., 1973; Lichter et al., 1997; Spitalny and Fenske, 1970; Sweeney et al., 2003).

Based on these phenotypic similarities, *Lmx1b*^*V265D*^ mutant mice are a valuable model for determining mechanisms and modifiers of ocular disease phenotypes in humans with *LMX1B* variants. Given the phenotypic variation between individuals with the same *LMX1B* variant (McIntosh et al., 2005), we expected to find differences in glaucoma-associated ocular phenotypes between different genetically diverse mouse strains with the *Lmx1b*^*V265D*^ allele. Here, we characterized the ocular effects of the *Lmx1b*^*V265D*^ allele on four different mouse strain backgrounds. Our results show that strain background significantly affects the onset and progression of glaucoma-related phenotypes in *Lmx1b*^*V265D/+*^ mice including IOP elevation and glaucomatous neurodegeneration. Based on this, we performed a gene mapping experiment between the most susceptible and resistant strain backgrounds and identified a modifier locus on Chromosome 18.

## Materials and Methods

### Animals husbandry and ethics statement

The *Lmx1b*^*V265D*^ mutation was discovered in an ENU mutagenesis screen (Cross et al., 2014; Thaung et al., 2002). It is formally named the *Icst* (iridocorneal strands) allele, but since it causes additional phenotypes and since other *Lmx1b* alleles cause iridocorneal strands we refer to it based on the protein level change V265D. Briefly, ENU-mutagenized Balb/cAnN (MRC Harwell, Oxfordshire, UK) were crossed to C3H/HeN mice (MRC Harwell, Oxfordshire, UK), and their offspring were screened (Thaung et al., 2002). Mice carrying the *Icst* mutation were crossed to C57BL/6J for gene mapping, and sequencing of the *Lmx1b* gene identified the *V265D* mutation (Cross et al., 2014; Thaung et al., 2002). These mice were then backcrossed to the C57BL/6J (Stock# 000664), DBA/2J-*Gpnmb*^*+*^*/*SjJ (Stock# 007048), C3A/BLiA-*Pde6b*^*+*^/J (Stock# 001912) and 129/Sj (Stock# 003884) for 8-10 generations). To determine if a tyrosinase deficient background has an earlier onset, more severe phenotype, *Lmx1b*^*V265D*^ was also backcrossed to the BALB/cJ (Stock# 000651) background for typically 6-8 generations and mice analyzed at 3 to 6 months of age. All experimental mice were backcrossed at least 6 generations. The 129/Sj strain was created from mice carrying a heterozygous knockout mutation generated in TL1 ES cells and maintained on a 129S6/SvEvTac background. We obtained this strain and selected mice without the heterozygous knockout mutation for inbreeding. Genotyping evidence suggests our 129/Sj strain contains large regions aligning to both 129S6/SvEvTac and 129S1/SvImJ respectively but is genetically distinct to any other 129 substrains (data not shown). DBA/2J, C3A/BLiA-*Pde6b*^*+*^/J, and 129/Sj mice were maintained on NIH 31 (6% fat) diet. To avoid obesity, C57BL/6J (B6) mice were maintained on NIH 31 diet (4% fat) diet and HCl acidified water (pH 2.8-3.2). Early studies showed that the minor difference in fat content did not affect the phenotypes. Mutant and control littermates were housed together with Alpha-dri bedding in cages covered with polyester filters. Cages were maintained in an environment kept at 21°C with a 14-hour light: 10-hour dark cycle. All mice were treated in accordance with the Association for Research in Vision and Ophthalmology’s statement on the use of animals in ophthalmic research. The Institutional Animal Care and Use Committee of The Jackson Laboratory approved all experimental protocols.

### Genotyping of the *Lmx1b* allele

*Lmx1b*^*V265D*^ and *Lmx1b*^*+*^ genotypes were determined using an allele-specific PCR protocol. Genomic DNA was PCR amplified with forward primer specific to the *V265D* allele 5’-TCAGCGTGCGTGTGGTCCTGGA-3’, a forward primer specific to the wild type allele 5’-GACATTGGCAGCAGAGACAGGCCGAGGCGTGCGTGTGGTCCATGT-3’, and the reverse primer 5’-ACACAAGCCTCTGCCTCCTT-3’. Genomic DNA was PCR amplified using the following program; 1) 95°C for 2 minutes, 2) 95°C for 15 seconds, 3) 57 °C for 20 seconds, 4) 72 °C for 30 seconds, 5) repeat steps 2-4 35 times, 6) 72°C for seven minutes. 5 μl of sample was run on a 3% agarose gel. The wild type allele amplifies a 175 base pair fragment and the *V265D* allele amplifies a 152 base pair fragment.

### Slit-lamp examination

Anterior eye tissues were examined approximately every 3 months between 2-13 months of age using a slit-lamp biomicroscope and photographed with a 40x objective lens. Phenotypic evaluation included iris structure, pupillary abnormalities, generalized corneal haze, corneal opacity, corneal keratopathy, hyphaema, hypopyon, corneal pyogenic granuloma, vascularized scarred cornea, buphthalmos, cataracts, and deepening of the anterior chamber. A subset of phenotypes that were common in *Lmx1b*^*V265D/+*^ mice (anterior chamber deepening, pupillary abnormalities, corneal haze, and corneal opacity) were characterized and graded based on a semiquantitative scale of either phenotype being not present, mild, moderate, or severe in presentation (Table 1). Detailed examination of typically 40 eyes from each strain and genotype at 4, 7, and 11 months of age was performed, except for the C3H background at 7 months where 12 mutant eyes and 14 WT eyes were examined. C3H mice at 4 months were examined but no phenotypes were graded. We found no sex difference in onset and severity of the phenotypes. Therefore, we combined both sexes in our analyses, with all cohorts including balanced numbers of male and female mice. Groups were compared pairwise by Fisher’s exact test.

**Table 1.**
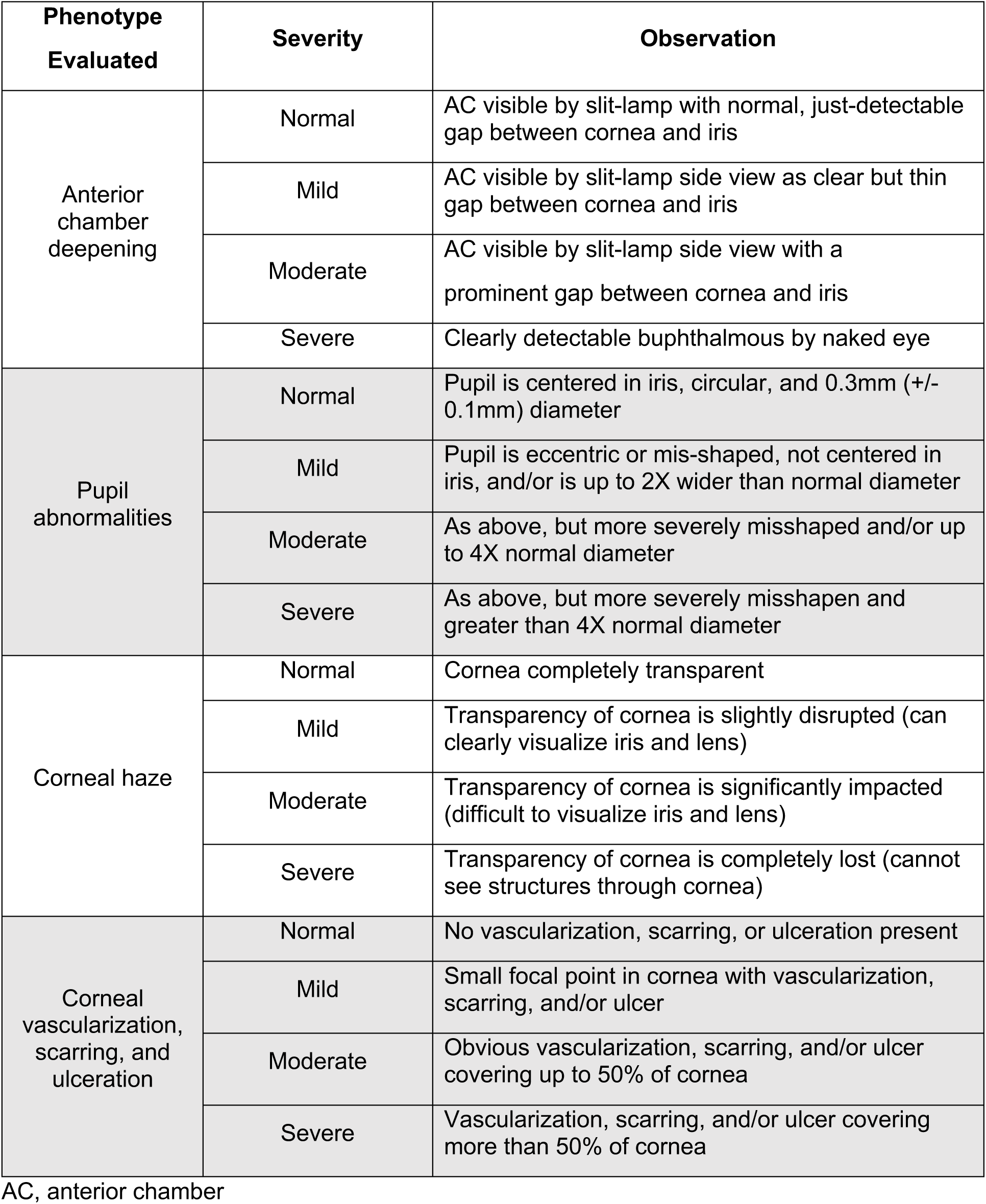
Severity definitions for abnormalities in *Lmx1b*^*V265D/+*^ mice.

### IOP measurement

IOP was measured using the microneedle method as previously described in detail (John et al., 1997; Savinova et al., 2001). Briefly, mice were acclimatized to the procedure room and anesthetized via an intraperitoneal injection of a mixture of ketamine (99 mg/kg; Ketlar, Parke-Davis, Paramus, NJ) and xylazine (9 mg/kg; Rompun, Phoenix Pharmaceutical, St. Joseph, MO) immediately prior to IOP assessment, a procedure that does not alter IOP in the experimental window (Savinova et al., 2001). IOP values were grouped by mouse age. IOPs measured at 3 to 5.9 months of age were grouped into the young timepoint (3-6mo), 6 to 8.9 months were grouped into the intermediate timepoint (6-9mo), and mice 9 to 11.9 months were grouped into the older timepoint (9-12mo). *Lmx1b*^*V265D/+*^ and WT IOP distributions did not meet the assumption of equal variance by Levene’s test. Therefore, we compared individual groups by two-tailed Welch’s *t*-test. In mice, IOP elevation caused by different mutations (including mutations in human glaucoma genes) is often accompanied by both an upward and a downward spread of values. This is due to complex effects including ocular stretching, perturbations of diurnal regulation and ciliary body dysfunction or atrophy (Chang et al., 2001; John et al., 1998). This spreading effect was strong in our current study and especially so for the *V265D* allele (likely exacerbated by their weakened/ expandable corneas and corneal ulceration with perforation in some mice - see text). Thus, to examine the magnitude of IOP dysregulation in *Lmx1b*^*V265D/+*^ eyes, we used the absolute value of the difference from the WT mean of each measurement (calculated by subtracting each mutant or WT value from the WT mean of the matching strain background and age). Distributions of these values were plotted and, as they also failed the assumption of equal variance between groups, were compared statistically by two-tailed Welch’s *t*-test. To further visualize the change in variance between *Lmx1b*^*V265D/+*^ and WT groups, we binned IOP values into four categories (<10mmHg, 10-19.9mmHg, 20-29.9 mmHg, and ≥30mmHg). The percentage of IOP values within each category was compared across experimental groups by Fisher’s exact test. We measured IOP of at least 30 eyes per group (age, genotype, and strain background) except for C3H background mice at 6-8 months where n= 20 mutant and 13 WT. All cohorts included balanced number of male and female mice. During each IOP measurement period, eyes of independent wild-type B6 mice were assessed in parallel with experimental mice as a methodological control to ensure proper calibration and equipment function.

### Ocular histological analysis

Enucleated eyes were fixed for plastic sectioning (0.8% paraformaldehyde and 1.2% glutaraldehyde in 0.08 M phosphate buffer (pH 7.4) as previously described in detail (John et al., 1998). Serial sagittal sections were collected, stained with hematoxylin and eosin, and analyzed for pathologic alterations at 3 months of age. For analysis of angle morphology relevant to drainage function, we used a previously validated grading scheme to determine the degree of angle closure due to adhesions/malformations that block drainage (Libby et al., 2003). The lower the total score the more extensively an angle is open around the circumference of any eye, while the higher the score the more closed it is. Briefly, we evaluated 24 similarly spaced angle regions from of each eye including the peripheral, mid-peripheral, and central ocular regions. For a few WT eyes, only 15 to 22 angle locations were scored due to regional processing artifacts. Angle scores for such eyes were normalized to the others for direct comparison. Each angle was graded based on the extent of angle blockage by attachment of the iris to the trabecular meshwork and cornea as previously reported (Libby et al., 2003) (0 = normal, iris and ciliary body join at iris root with no adhesion to the TM or cornea, 1 = iris attached to very posterior portion of TM so that most of the TM/angle is open and accessible for drainage, 2 = iris attached to TM for up to three quarters of the extent of TM, 3 = iris covers entire TM and extends just into peripheral cornea indicating a completely closed angle region, 4 = iris covers TM and adhesion extends further onto cornea). The final angle score is the sum of values for each angle location. The minimum possible score is 0, reflecting a completely normal, fully-open angle at all locations. A score of 24, would indicate that either the angle is completely open for at least 75% of it assessed circumference with minor abnormalities in the remaining 20%, or that an angle is open for even more of its circumference with focal occurrence or more severe abnormalities. The maximum score is 96 (4 × 24) reflects a completely closed angle at all locations with extensive attachment of the iris to the peripheral cornea. A score of 72 (3 × 24) also reflects a completely closed angle as the iris completely covers the TM at all assessed locations in such eyes. The samples were intermixed, and the observers were not aware of the *Lmx1b* genotype or genetic background during the grading. Two observers masked to sample identity as well as themselves graded the eyes. The score assigned by each observer agreed >96% of the time and never disagreed by more than 1 grade. Disagreements involved regions with abnormalities at the border of two grades and differences were resolved by consensus agreement when still masked to sample identity. The summed grade of all the examined angles from each mouse is plotted. Because the data was discretized, groups were statistically compared by Mann-Whitney U test. Each group contained balanced numbers of male and female mice. We analyzed 5-7 eyes, with a median of 6 eyes of each genotype for both the 129 and B6 strains.

### Optic nerve assessment

Intracranial portions of optic nerves were dissected, processed, and analyzed as previously described (Howell et al., 2007; Howell et al., 2012; Nair et al., 2016; Williams et al., 2017). Briefly, optic nerve cross-sections were stained with para-phenylenediamine (PPD) and examined for glaucomatous damage. PPD stains all myelin sheaths of a healthy axon, but differentially darkly stains the myelin sheaths and the axoplasm of sick or dying axons. This allows for the sensitive detection and quantification of axon damage and loss. Optic nerves were prepared for analysis with a 48h fixation in 0.8% paraformaldehyde and 1.2% glutaraldehyde in 0.08M phosphate buffer (pH 7.4) at 4°C followed by overnight treatment in osmium tetroxide at 4°C. Nerves were washed twice for 10 minutes on 0.1 M phosphate buffer, once in 0.1 M sodium-acetate buffer and dehydrated in graded ethanol concentrations. Tissues were then embedded in Embed 812 resin (Electron Microscopy Sciences, Ft. Washington, PA), and 1-μm thick sections were stained in 1% PPD for approximately 40 minutes. Stained sections were compared using a previously reported grading scale that is validated against axon counting (Howell et al., 2007; Howell et al., 2012). All cohorts included balanced numbers of male and female mice. We analyzed approximately 30 nerves for each strain and genotype, except for strain 129 WT and D2-G mutant groups where we graded 15 and 17 nerves per group respectively. Groups were compared pairwise by Fisher’s exact test.

### Gene mapping and QTL analysis

To identify loci controlling strain differences in phenotype onset and severity, *Lmx1b*^*+/+*^ males of the glaucoma-susceptible B6 background were crossed to glaucoma-resistant 129.*Lmx1b*^*V265D/+*^ female mice. 129B6F1 *Lmx1b*^*V265D/+*^ mice were screened for anterior eye phenotypes by slit-lamp between 1-6 months. 129B6F1 mice were resistant to *Lmx1b’s* effects, indicating that a dominant 129 locus(i) contributes to phenotypic resistance. To characterize this locus(i), we backcrossed 129B6F1 *Lmx1b*^*V265D/+*^ mice of both sexes to the B6 background to create an N2 recombinant mapping cohort. A total of 107 N2 *Lmx1b*^*V265D*/+^ progeny of both sexes were aged and screened by slit-lamp and IOP measurement between 1-3 months and 4-5 months. We used slit-lamp data to map the genomic loci contributing to resistance. Based on slit-lamp data, each mapping mouse was binned into one of three categories; *bilateral susceptible* (B6-like), *bilateral resistant* (129-like), or *unilateral*. Mice were considered *bilateral susceptible* if one or more of the following phenotypes was at least moderate or severe in each eye; anterior chamber depth, pupil open, corneal haze, corneal opacity. If neither eye had any moderate or severe ocular phenotypes, mice were considered *bilateral resistant*. Due to variable expressivity several mice had *unilateral phenotypes affecting only the left or right eye in a random fashion*. Genotyping was performed using 138 regularly spaced genome-wide single nucleotide polymorphic markers that differentiate the B6 and 129 genomes (SNPs, KBioscience, UK). We performed a genome-wide one-dimensional quantitative trait locus (QTL) scan to identify the chromosomal loci modulating *Lmx1b* phenotypes. r/QTL version 1.14-2 was used for QTL analysis (Broman et al., 2003). The final QTL analysis uses mouse sex as an additive covariate and calculates genotype probability between SNP markers. QTL intervals were based on a 1.5 LOD drop from the maximum LOD peak on the chromosome. Because the first marker we genotyped on Chr 18 was at 5 Mb, we did not examine recombinants between 0-5 Mb. All genomic coordinates were calculated using GRCm38 (mm10) assembly.

### Congenic strain generation and phenotyping

The implicated, strain 129, Chr 18 locus was then backcrossed to B6. At each generation of backcrossing, we used 4 microsatellite (MIT) markers to genotype the 69.5 Mb Chr 18 interval (D18MIT88, D18MIT123, D18MIT185, and D18MIT19). After 10 or more generations, mice heterozygous (B6/129) at each of these Chr18 markers were crossed to B6.*Lmx1b*^*V265D*/+^ mice to generate our experimental cohort. Chr 18 was genotyped in each experimental mouse using the same 4 MIT markers. Mice were assessed by slit-lamp at 1-6 months of age and binned using the same *bilateral susceptible, bilateral resistant*, and *unilateral* groupings as for the N2 mapping mice. All groups contained balanced numbers of males and females and were compared by Fisher’s exact test.

## Results

### Strain 129 background is most resistant while B6 background is most susceptible

Strain background had a profound effect on the ocular phenotypes in *Lmx1b*^*V265D/+*^ mice (Fig. 1). Rare abnormal phenotypes were detected in some WT mice. This is due to the previously documented susceptibility of B6 mice to developmental abnormalities including anterior segment dysgenesis and anophthalmia (Chase, 1942; Gould and John, 2002; Smith et al., 1994). The frequency of these abnormalities varies based on factors such as environmental stress, and alcohol exposure (Cook et al., 1987; Sulik et al., 1981; Webster et al., 1983). Ocular disease phenotypes in *Lmx1b*^*V265D/+*^ mutants included deepened anterior chambers, malformed and eccentric pupils, iridocorneal strands (strands of iris focally fused to cornea), corneal haze, corneal vascularization, corneal scleralization, and corneal ulceration (Fig. 1). We compared group differences in the frequency and severity of ocular phenotypes using Fisher’s exact test. Of all examined strain backgrounds, 129.*Lmx1b*^*V265D/+*^ mice were most resistant to developing these abnormal ocular phenotypes (Fig. 1). When present in strain 129 mutants, phenotypes were generally mild (Figs 1,2). Overall, B6 mutants had the most developmentally severe phenotypic abnormalities of all backgrounds. Compared to D2-G mutants, B6 mutants develop more severe anterior chamber deepening at young ages (3-5 months) and more severe corneal haze at all ages (all *P* < 0.01, Fig. 2). C3H and B6 mutants were similar in phenotype severity across ages, except for corneal haze, which was significantly more severe in B6 mutants at young and intermediate ages (6-8 months; *P* < 0.01, Fig. 2). Therefore, overall B6.*Lmx1b*^*V265D/+*^ are the most susceptible, C3H and D2-G backgrounds are intermediate, while strain 129 is the most resistant.

**Figure 1:**
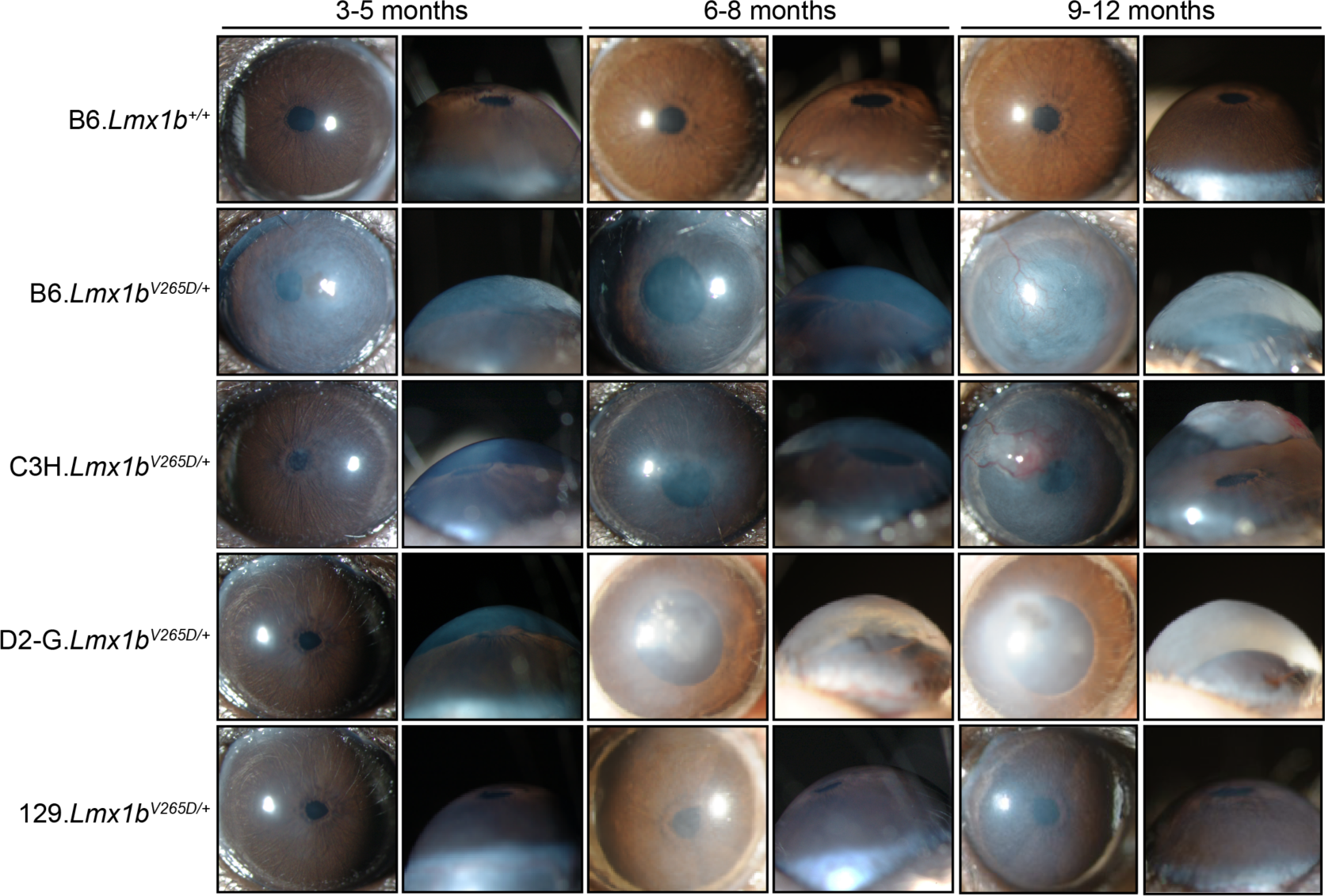
Strain background alters phenotypes in *Lmx1b*^*V265D/+*^ mice. Representative front and side-view, slit-lamp images for mice of the indicated ages and genotypes. The frequencies of specific disease features are shown in Figure 2. WT mice of all backgrounds were similar and so only B6 WTs are shown.B6.*Lmx1b*^*V265D/+*^ mutant mice have the most severe overall phenotypes including malformed eccentric pupils, extensive corneal haze and greatly deepened anterior chambers at 3 months of age. With age, the severity of B6 phenotypes increases, with development of corneal scarring, vascularization, and ulcers. C3H mutants are generally similar to B6 but are more resistant to developmental corneal phenotypes at younger ages. D2-G mutants are generally similar to C3H, but more resistant to LMX1B-induced corneal phenotypes at all ages (see Figure 2). The 129 strain background is the most resistant, with mutants typically displaying only mild pupillary abnormalities and corneal haze.

**Figure 2:**
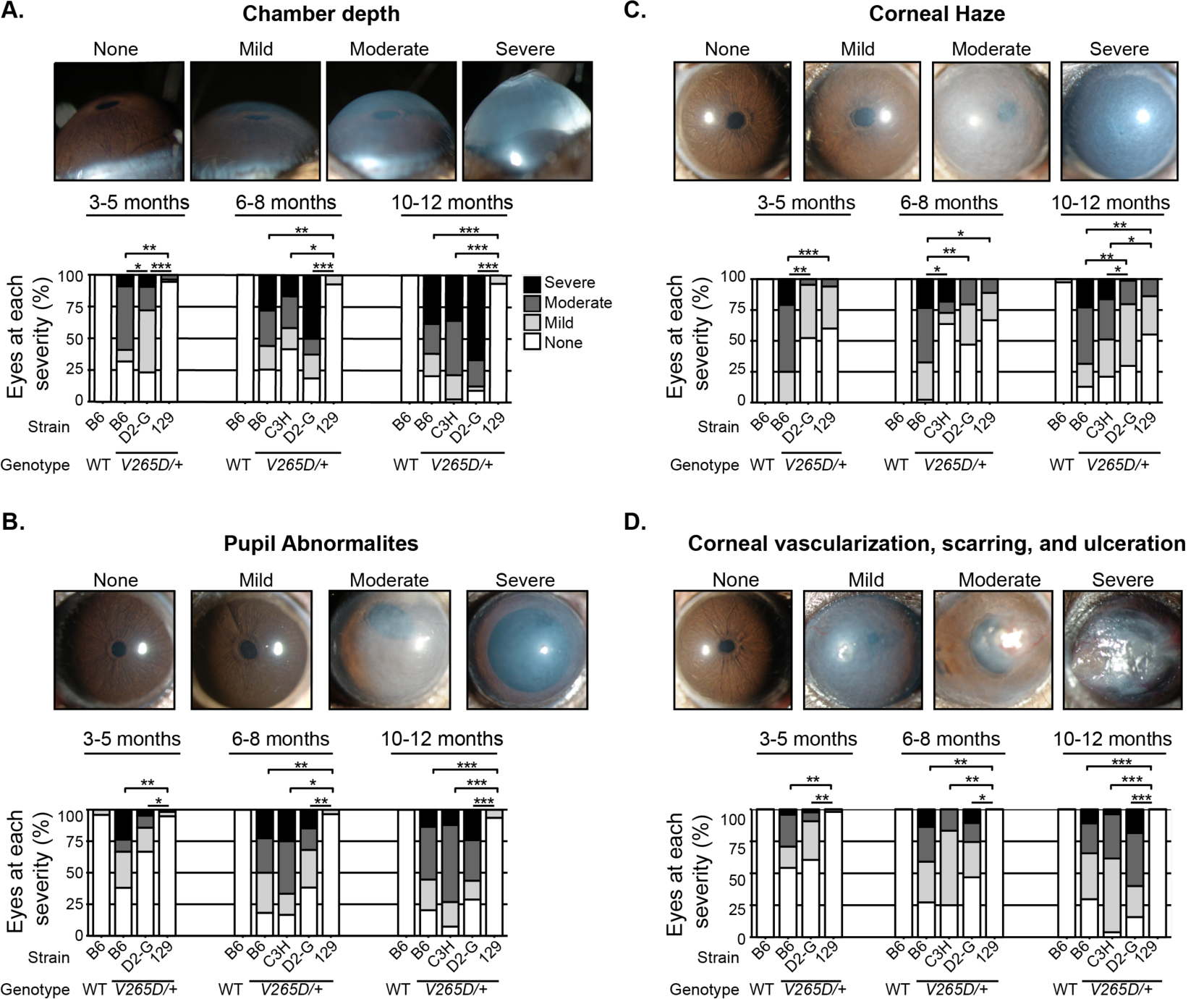
B6 background is most susceptible, while strain 129 is most resistant. **(A)** At 3-5 months, B6 mice have the most severe anterior chamber deepening (ACD), even compared to D2-G mutants (*P* = 0.0034). Strain 129 mutants rarely develop abnormal ACD at any age. Anterior chamber deepening (ACD) is a symptom of IOP elevation. (B-D) The same was true for corneal haze, pupillary and corneal abnormalities. * *P*< 0.01; ** *P* < 1.0E-05; *** *P* < 1.0E-10 (see supplementary table 1 for exact *P* values).

To test if the phenotypic differences are impacted by strain-dependent functional changes in the WT *Lmx1b* locus, we examined the *Lmx1b* locus of all four inbred strains using sequence data available from the Sanger Mouse Genomes Project (Keane et al., 2011). As 129/Sj and C3A/BLiA-*Pde6b*^*+*^/J were not available in the database, we used three closely related substrains of 129 (129P2/OlaHsd, 129S1/SvlmJ, and 129S5SvEvBrd) and the closely related C3H/HeJ substrain respectively as proxies for the strain 129 and C3H genotypes. Compared to the B6 reference genome, neither the 129 nor C3H substrains have any coding regions or intergenic variants in conserved regions that would affect function. In contrast, the D2 background contains 3’ UTR variants, synonymous coding variants, and a predicted splice region variant (rs27178126) 8bp from the splice donor site in intron 3. However, transcriptomic data from D2-G background limbal tissue showed no splicing abnormalities of any *Lmx1b* exons (data not shown). Given this, and the facts that 1) WT D2 mice lack haploinsufficient *Lmx1b* phenotypes and 2) *Lmx1b*^*V265D/+*^ D2-G mice are phenotypically similar to heterozygotes on the others strains and lack lethal homozygous mutant phenotypes, we conclude that the splice-region change has no effect. Additionally, as our most susceptible and resistant strains have identical *Lmx1b* loci, there is no clear relationship between the strain-specific, WT, *Lmx1b* locus and phenotypic severity. This indicates that other genetic modifier(s) underlie the observed phenotypic differences between these strains.

### B6.*Lmx1b*^*V265D/+*^ mice have the most severely affected drainage structures

Structural abnormalities in the aqueous humor drainage structures, Schlemm’s canal (SC) and trabecular meshwork (TM), can lead to glaucoma by impacting IOP. These structures are located within the iridocorneal angle that runs around the entire limbal circumference of the eye. To evaluate whether strain background impacted drainage structure abnormalities in *Lmx1b*^*V265D/+*^ mice, we analyzed the morphology of the iridocorneal angle of our most extreme strains B6 and 129. WT mice have open drainage angles and normal SC and TM morphology (Fig. 3A). We found a spectrum of abnormalities in *Lmx1b*^*V265D/+*^ mice of both B6 and 129 backgrounds including malformed or absent SC and/or TM as well as iridocorneal angle adhesions causing an obstructed aqueous humor drainage route (closed angle; Fig. 3A). The severity of abnormalities in mutant mice varies both between eyes and locally around the angle circumference within individual eyes (Fig. 3A). Importantly, angle abnormalities were more severe in B6 background mutants compared to those of the 129 background (Mann-Whitney U Test, *P* = 0.0023; Fig. 3B). Despite open-angle regions, B6 mutant angles were closed to aqueous humor drainage around much of the ocular circumference. However, strain 129 mutant angles were largely open and typically had only mild abnormalities. Mild iridocorneal angle abnormalities have been reported in patients with *LMX1B* variants and POAG (Lichter et al., 1997; Vollrath et al., 1998)

**Figure 3:**
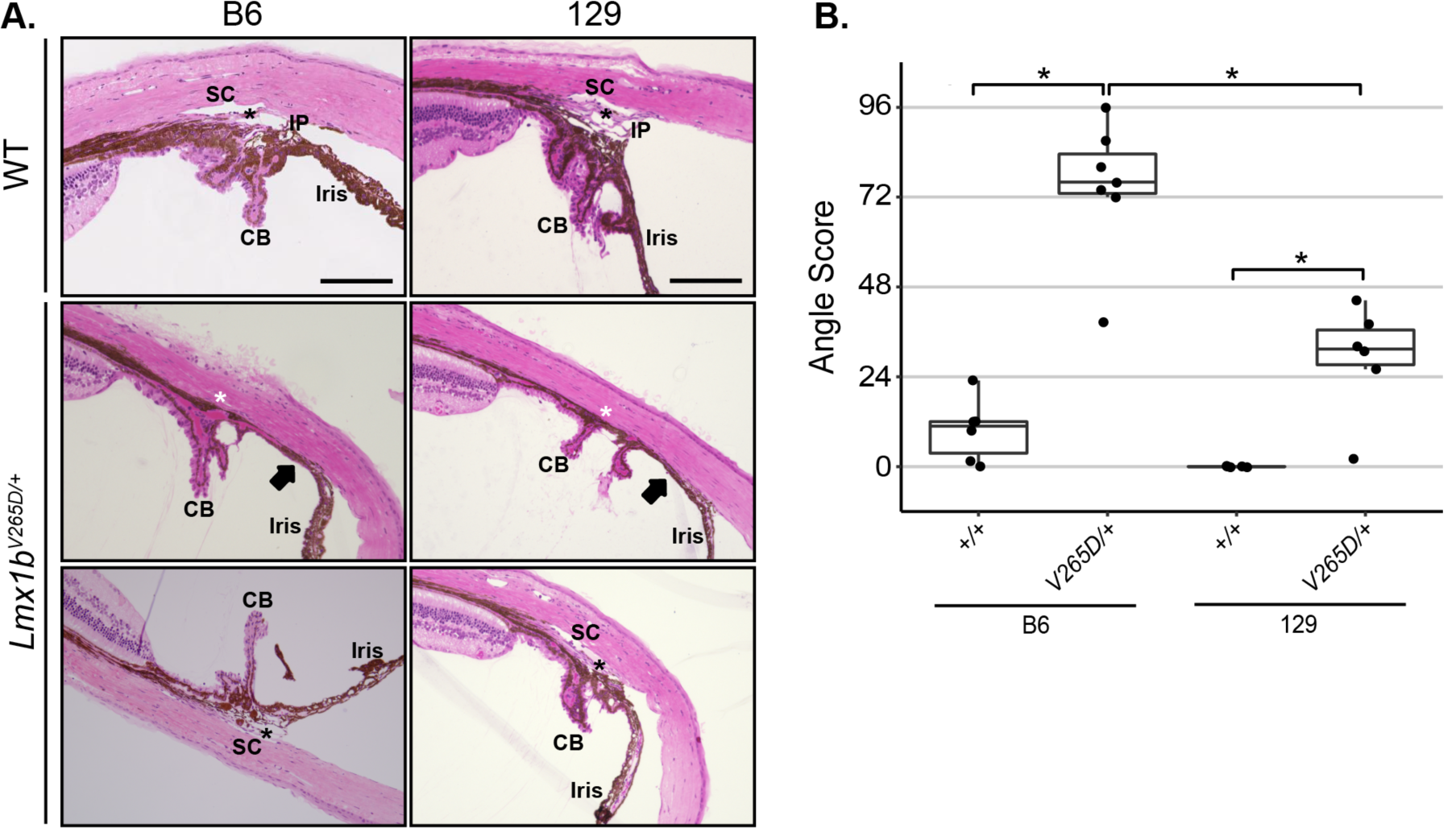
129.*Lmx1b*^*V265D/+*^ mutants are resistant to angle abnormalities compared to B6. **(A)** Representative images of the iridocorneal angle region (H&E stained sagittal sections). WT mice of both B6 and 129 backgrounds have normal open-angle morphology. Narrow iris processes (IP), are known to occur intermittently around the angle of WT mice without obstructing aqueous humor drainage. SC: Schlemm’s canal, black asterisk: trabecular meshwork, CB: ciliary body. In mutant eyes, abnormalities, including severe iridocorneal adhesions (arrows) as well as absent (white asterisk) or hypomorphic SC and TM, are locally present within individual eyes (middle panels) with different locations within the same eyes having open-angles of normal morphology (bottom panels). Scale bar = 200μm. **(B)** B6 mutant have high angle scores (Methods), indicating largely closed or malformed angles. Strain 129 mutants have less severely affected, largely open-angles. Higher angle scores indicate a more severely and more extensively affected angle around its circumference. A score of 96 represents a severely abnormal angle at all locations while an angle with a score of 0 being is completely normal and open at all locations. The strain 129 median grade of 31 indicates that their angles were open at most locations around the eye. It is established that a small incidence of developmental abnormalities occurs in B6 WT mice (see main text). Boxplots show interquartile range and median line. Mann-Whitney U test; * = *P* < 0.01 (129 vs B6 mutants, *P* = 0.0023; 129 WT vs mutant *P* = 0.0055; B6 WT vs mutant, *P* = 0.0033). All eyes were examined at 3-months-old. Between 5 and 7 eyes examined for each genotype and strain.

### IOP is abnormal in *Lmx1b*^*V265D/+*^ mice of all backgrounds with B6 being most severe at young ages

We longitudinally examined IOP in WT and *Lmx1b*^*V265D/+*^ eyes and found an overall change to the distribution of IOP values in *Lmx1b*^*V265D/+*^ eyes compared to WT controls. Spreading of IOP in both directions can be caused by various reasons including ciliary body atrophy/malformation and corneal damage, as is most common in the *Lmx1b* B6 mutants here. Across all strain backgrounds and ages, mutant eyes had both the highest and lowest IOP values (Fig. 4A-C). The variance of WT and mutant IOP distributions was significantly different at all examined ages in B6, C3H, and D2-G backgrounds (Levene’s test, all *P* < 0.01). We found significantly elevated IOP in C3H (6-9 months), D2-G (3-6 and 6-9 months), and strain 129 (3-6 and 6-9 months) mutants compared to WT controls (Fig. 4A,B; Welch’s *t*-test, all *P <* 0.01). Although IOP was significantly elevated, the spreading of values in both directions masked the ability to detect mean differences compared to WT controls for other mutant groups (see methods). At 3-6 months, B6 mutants have a larger average dispersion (absolute difference from WT mean) than D2-G and strain 129 mutants (Welch’s *t*-test, all *P* < 0.01; Fig. 4D). Consistent with this, 10% of B6 mutant eyes had IOP >30mmHg at 3-6 months, a magnitude not found in age-matched WT or mutant eyes of any other backgrounds (Fig. S2). C3H mutants did not have IOP assessed during the 3-6 months age window but appeared similar to B6 in anterior chamber deepening, a reflection of raised IOP. Although IOP abnormalities were detected in strain 129 mutants at different ages, there was significantly less IOP dispersion in these mice at advanced ages (9-12 months) compared to all other backgrounds (Welch’s *t*-test, all *P* < 0.01; Fig. 4F). Overall, our data shows that *Lmx1b*^*V265D*^ has a strong impact on IOP with the most extreme and earliest phenotypes on a B6 background.

**Figure 4:**
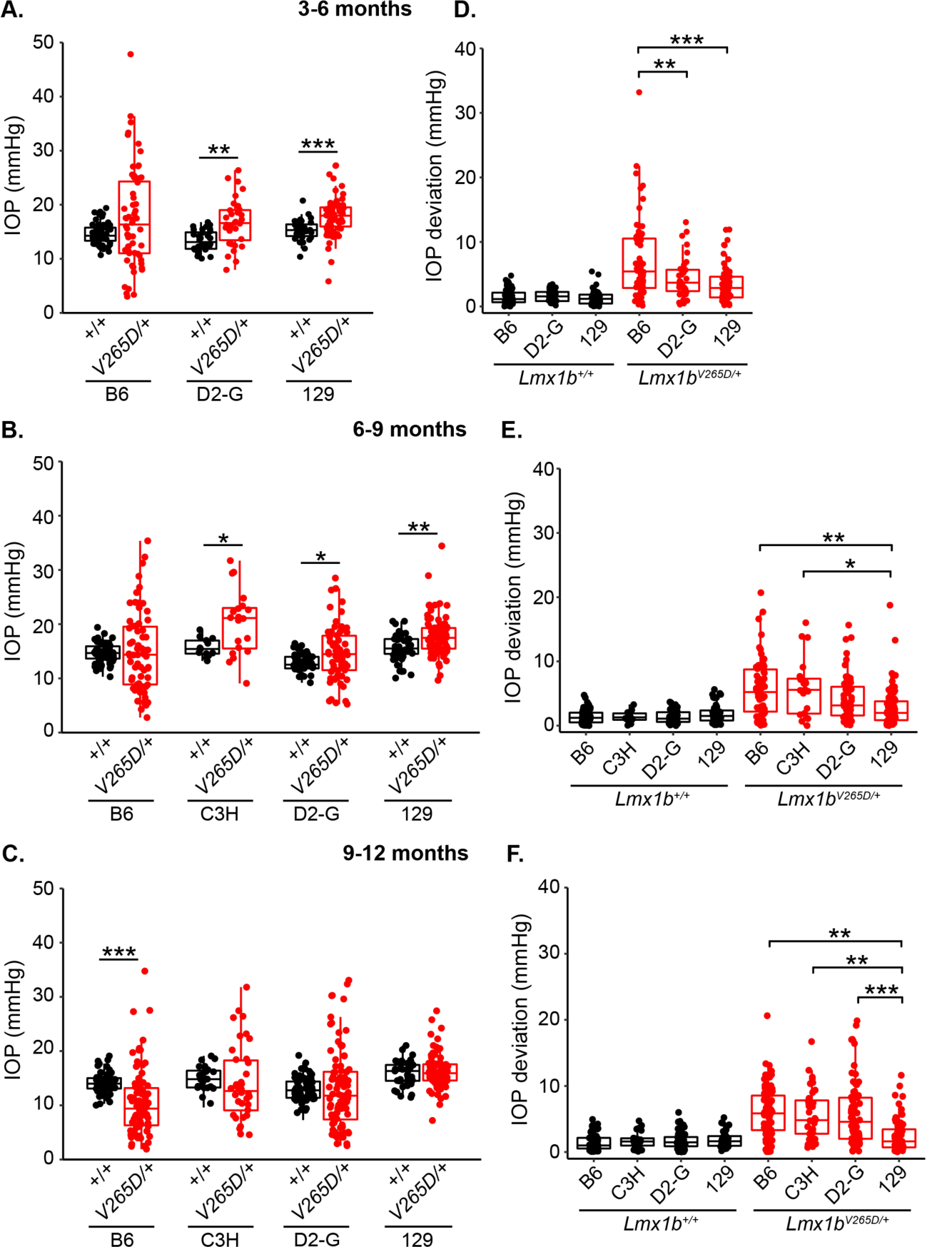
IOP in *Lmx1b* mutants. **(A-C)** Boxplots of IOP (interquartile range and median line) clearly indicate spreading of IOP in mutants of all strain backgrounds with clear IOP elevation in some mutants. **(A-B)** *Lmx1b*^*V265D/+*^ mutants of D2-G and strain 129 backgrounds have significantly elevated IOP compared to respective WT controls at 3-6 mo and 6-9 mo. C3H mutants have elevated IOP at 6-9 months compared to WT controls (*P* = 0.0032). Although IOP was not measured, anterior chamber deepening suggests IOP is elevated in many C3H mutants prior to 6 months age (Figure 1) **(C)** Due to an increase in abnormally low IOP values, B6.*Lmx1b*^*V265D/+*^ mice have a significantly lower IOP average than WT controls at 9-11 months old (*P* = 8.2E-7). **(D-F)** Boxplots of IOP deviation (absolute value of difference to respective WT mean value, Methods) At all ages, WT groups had minimal IOP deviation, with no values deviating more than 7mmHg. **(D)** At 3-5 months, B6 mutants have a significantly greater IOP deviation compared to those of D2-G (*P* = 0.0068) and strain 129 (*P* = 4.5E-05) backgrounds. **(E-F)** Strain 129 mutants have significantly less IOP deviation compared to B6 and C3H mutants at 6-9 months and to all other backgrounds 9-12 months. * *P* < 0.01, ** *P* < 0.001, *** *P* < 0.0001 (see supplementary table 2 for exact *P* values).

### B6.*Lmx1b*^*V265D/+*^ mice develop severe glaucoma but 129.*Lmx1b*^*V265D/+*^ mice do not

To assess the extent to which IOP elevation leads to glaucomatous neurodegeneration across genetic backgrounds, we histologically assessed retinas and optic nerves of *Lmx1b*^*V265D/+*^ and WT mice. Because the majority of abnormally elevated IOP values are found at 3-6 and 6-9 months in B6.*Lmx1b*^*V265D/+*^ mice, we examined their optic nerves between 10-12 months. Optic nerves of D2-G and 129 backgrounds were examined slightly later in life (12-14 months). Consistent with other ocular phenotypes, B6.*Lmx1b*^*V265D/+*^ mice had the highest prevalence of severely degenerated optic nerves with nearly 80% of nerves having severe axon loss and damage and prominent gliosis (Fig. 5A,B). Importantly, mutants on the 129 background did not develop any detectable optic nerve degeneration (Fig. 5A), even at the oldest age examined. Mutants with optic nerve degeneration had characteristic hallmarks of glaucoma with retinal nerve fiber layer thinning (layer containing retinal ganglion cell axons) and optic nerve excavation/remodeling (Fig. 5C).

**Figure 5:**
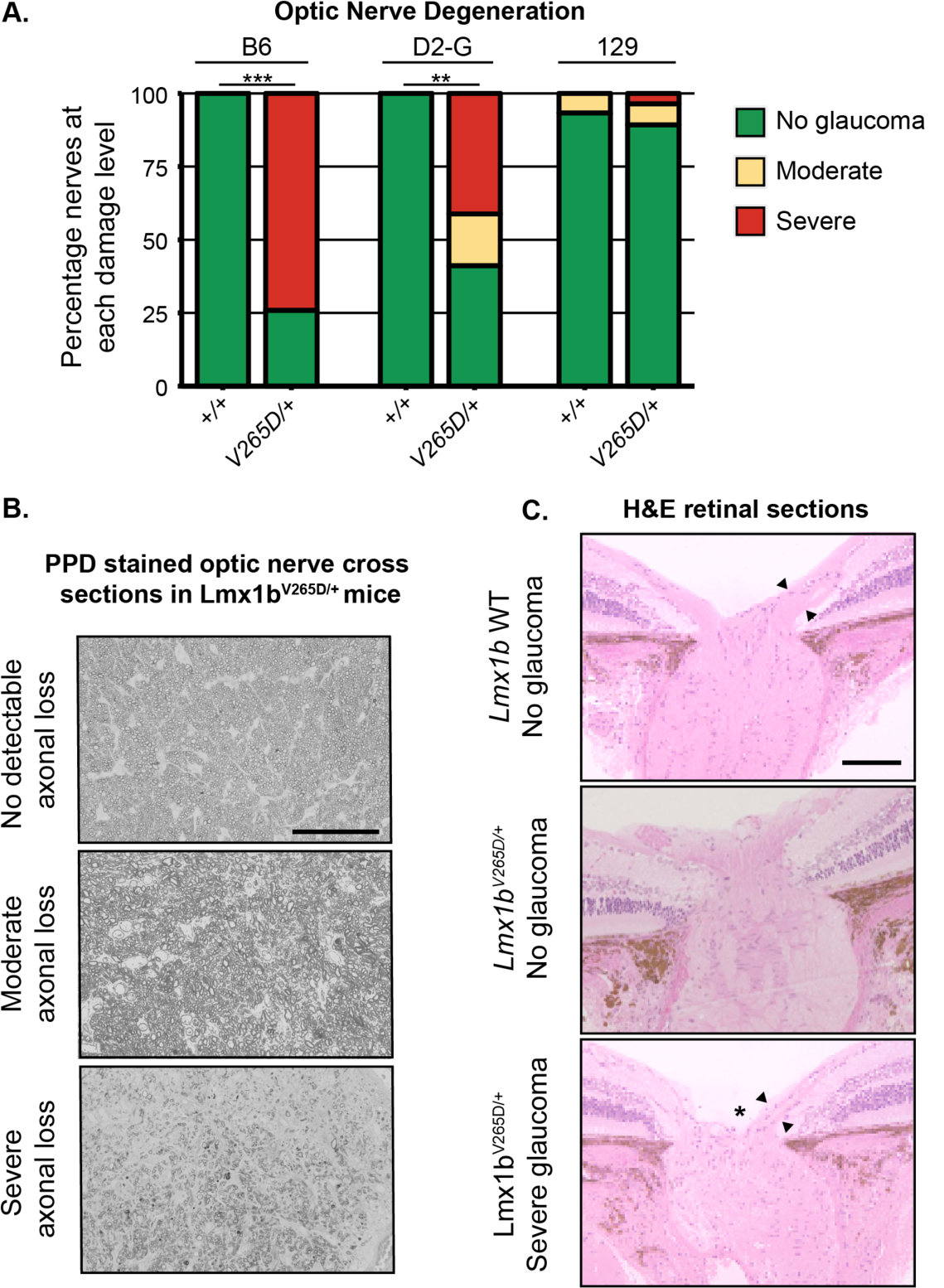
B6.*Lmx1b*^*V265D/+*^ mice develop glaucomatous neurodegeneration while 129.*Lmx1b*^*V265D/+*^ mice do not. **(A)** Frequency histogram of degree of optic nerve damage evident in PPD stained cross sections (Methods) ** *P* < 1.0E-05, *** *P* < 1.0E-10. **(B)** Representative images of PPD-stained optic nerve cross sections from *Lmx1b*^*V265D/+*^ mice. (Top) Healthy nerves at 10 months old had no detectable axonal damage. These axons had a clear axoplasm and darkly stained myelin sheaths. (Middle) Moderate optic nerve degeneration with some axon loss and early gliosis. (Bottom) Severe damage and extreme axon loss with extensive glial scarring. Scale bar = 50μm **(C)** H&E stained optic nerve heads with flanking retina. WT eyes have normal nerve heads with a thick nerve fiber layer (arrowheads) as do unaffected mutants. Severely affected mutants have pronounced optic nerve excavation (asterisk) with loss of the nerve fiber layer (arrowheads), Scale bar = 200μm.

### The BALB background does not increase disease severity

Mutation of the tyrosinase gene (*Tyr*) increases susceptibility to ocular drainage tissue defects in *Cyp1b1* and *Foxc1* mutants (Libby et al., 2003). To assess the effect of a further genetic background and if *Lmx1b-*induced disease onset is earlier in a *Tyr* deficient background, we crossed the *Lmx1b*^*V265D/+*^ allele to the albino BALB/cJ (BALB) strain background and analyzed 3 to 6 months old mice (Yokoyama et al., 1990). Compared to our most susceptible B6 background, BALB mutant mice are resistant to *Lmx1b*-induced anterior segment developmental phenotypes (Fig. S3, Table S1). Similar to strain 129 mutants, anterior segment phenotypes in BALB mutants were generally mild when present. BALB mutants did develop elevated IOP with their IOP distribution being similar to the other backgrounds (Fig. S3, Table S2). Therefore, *Tyr* genotype did not exacerbate disease severity in *Lmx1b* mutants on a BALB background.

### A locus on Chromosome 18 determines differential susceptibility to *Lmx1b*-associated phenotypes

In order to identify genomic regions contributing to differential susceptibility between the B6 and 129 genetic background, we performed a mapping cross. Specifically, F1 progeny were created using males from the susceptible B6 background and *Lmx1b*^*V265D/+*^ females from the more resistant 129 background. 129B6 *Lmx1b*^*V265D/+*^ F1’s phenocopied 129 mutants indicating 129-dominant loci confer disease resistance (data not shown). Thus, we backcrossed F1’s to B6 to generate our N2 mutant mapping progeny. Based on the severity of ocular phenotypes at 2 months of age, N2 *Lmx1b*^*V265D/+*^ mice were binned into a *bilateral susceptible* (B6-like), *bilateral resistant* (129-like), or *unilateral* categories. The unilateral category was used when only a single eye displays severe abnormalities and reflects reduced susceptibility to *Lmx1b*^*V265D/+*^-induced phenotypes. All N2 mapping progeny were genotyped using single nucleotide polymorphic (SNP) marker analysis. Using this data, we performed a quantitative trait locus (QTL) scan in our N2 cohort against ocular phenotype severity. In 1 to 3 months old mice, we detected intervals on Chromosomes 1 (33-139 Mb, max LOD at 53.6 Mb) and 18 (5-71.7 Mb, max LOD at 30.9 Mb) that significantly associated with the mapping phenotypes using a genome-wide significance cutoff (Fig. 6A). At 4 to 5 months, however, only the interval on chromosome 18 (5-74.5 Mb, max LOD at 30.9 Mb) reached genome-wide significance (Fig. 6B). to test whether this locus is sufficient to generate resistance to disease phenotypes in *Lmx1b* mutants, we backcrossed the strain 129 Chr 18 interval onto the B6 background. B6.*Lmx1b*^*V265D/+*^ mice that were heterozygous B6/129 throughout the Chr 18 interval were significantly more resistant to the *Lmx1b*-induced phenotypes than littermates that were homozygous B6 (Fisher’s exact test, P=0.0036; Fig. 6C). This validates this resistance locus.

**Figure 6:**
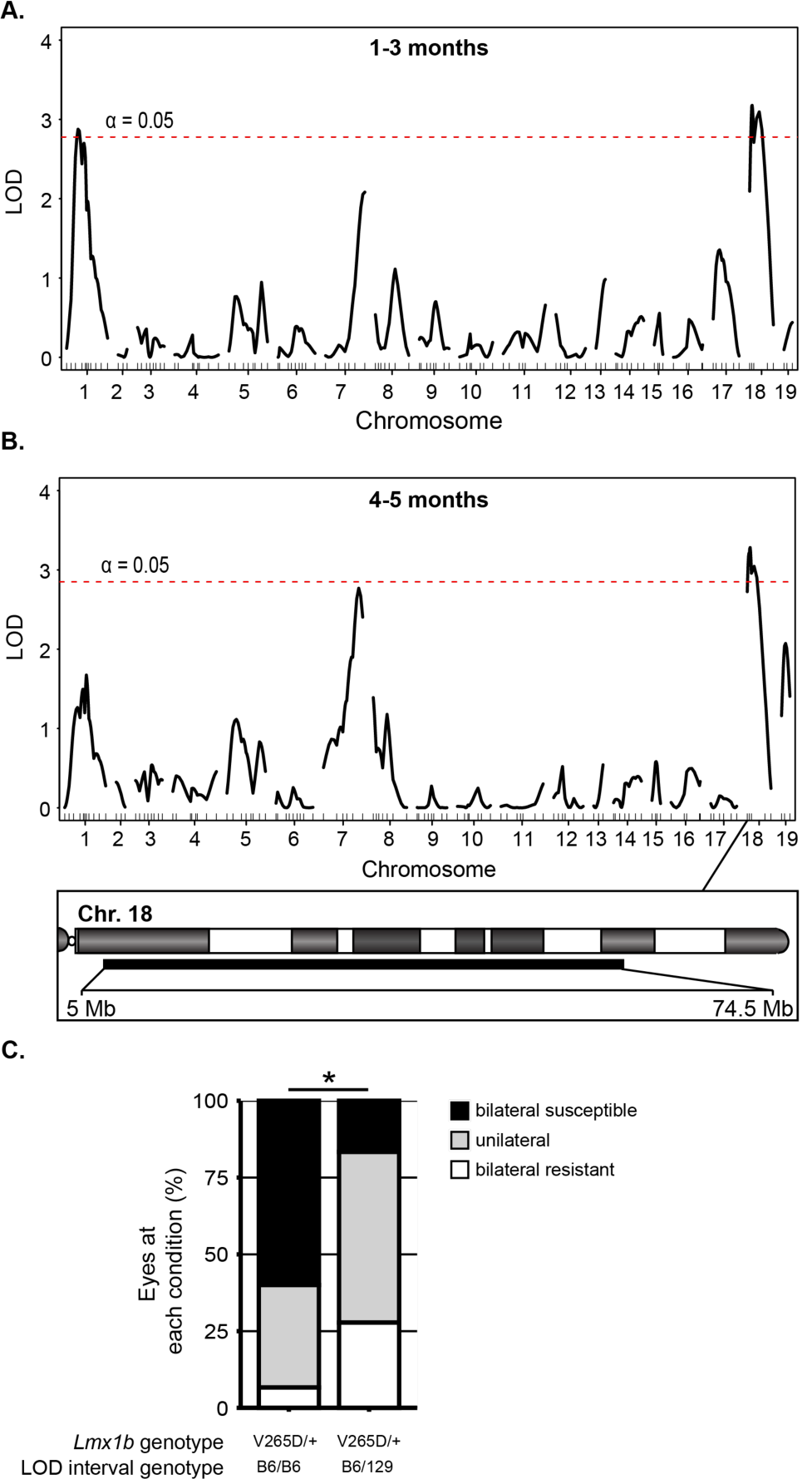
Modifier loci. Based on slit-lamp data, individual N2 mice were binned into one of three categories; *bilateral susceptible* (B6-like), *bilateral resistant* (129-like), or *unilateral*. Using this data, a genome-wide one-dimensional quantitative trait locus (QTL) scan was performed. **(A)** At 1-3 months, intervals on both Chr 1 (33-139 Mb, max LOD at 53.6 Mb) and Chr 18 (5-71.7 Mb, max LOD at 30.9 Mb) reached genome wide significance (5% significance threshold, genome-wide corrected, red dotted line). **(B)** At 4-5 months, an interval on Chr18 (5-74.5 Mb, max LOD at 30.9 Mb) with the same max LOD as 1-3 months was identified at genome wide significance. **(C)** Testing of the modifier locus by comparing *Lmx1b*^*V265D/+*^ mutant mice that are either homozygous (B6/B6) or heterozygous (B6/129) for the Chr 18 intervals. Having a strain 129 genotype throughout the modifier interval significantly increased resistance to severe ocular phenotypes compared to B6 homozygous littermates (Fisher’s exact test, *P* = 0.036). *Lmx1b* WT mice that are B6/129 heterozygous for the Chr 18 interval did not develop anterior eye phenotypes (data not shown). At least 15 mice examined per group.

## Discussion

### Differing disease presentation between individuals

Recent GWAS studies indicate that *LMX1B* variants cause elevated IOP and glaucoma in the general human population, without evident anterior segment abnormalities, involvement of other organs/tissues or NPS diagnosis (Choquet et al., 2018; Gao et al., 2018; Gharahkhani et al., 2018; Khawaja et al., 2018; MacGregor et al., 2018; Shiga et al., 2018). Similarly, *LMX1B* mutations cause organ specific kidney disease without extrarenal involvement (Boyer et al., 2013; Isojima et al., 2014). Several factors may contribute to differing disease presentations between individuals including the nature of the *LMX1B* variant, genetic modifiers, and environmental factors. Here, we clearly show that genetic background has a strong influence on disease presentation. This effect of genetic background allows a path to deciphering key pathogenic mechanism through characterization of modifier genes. Additionally, this effect must be considered when interpreting experimental data. For example, previous studies report that mice heterozygous for a null allele of *Lmx1b* have normal eyes on both a C57BL/6J and a C57BL/6×129/Sv mixed background respectively (Cross et al., 2014; Pressman et al., 2000). Haploinsufficiency is generally accepted to contribute to human disease as heterozygous deletions including *LMX1B* are pathogenic (Bongers et al., 2008; McIntosh et al., 1998). Thus, it remains unclear if mice differ to humans in their sensitivity to haploinsufficiency-induced phenotypes or if null alleles will induce characteristic abnormalities when assessed on further genetic backgrounds.

The nature of the mutation in *Lmx1b* is important to consider. The pathogenic nature of *LMX1B* haploinsufficiency suggests reduced transcription factor dosage or activity causes disease. However, as demonstrated by the *Lmx1b*^*V265D*^ allele, different mechanisms apart from haploinsufficiency can contribute to glaucoma such as dominant negative effects (Cross et al., 2014). The location of the point mutation within human *LMX1B* correlates with disease severity in the kidney (Bongers et al., 2005). Additional functional characterization of *LMX1B* mutations is required to better understand how the nature of the *LMX1B* variant affects disease onset and severity. Recently, a dominant stop codon mutation (*Lmx1b*^*Q105X*^, reported as *Lmx1b*^*Q82X*^) was shown to cause IOP elevation and glaucoma without anterior segment developmental abnormalities (by slit-lamp) on the D2-G background (Choquet et al., 2018). This contrasts to the *Lmx1b*^*V265D*^ allele, which induces obvious anterior segment abnormalities on the same D2-G genetic background (Figs 1-3). Together, these data strengthen the suggestion that the nature of individual *LMX1B* alleles affects the range and severity of disease outcomes in human patients (Bongers et al., 2005; McIntosh et al., 1998).Characterizing different mutant alleles on genetically diverse backgrounds will be important in determining disease mechanisms and discovering genetic modifiers, with the goal of improving risk assessment and developing therapeutics (Jeanne and Gould, 2017).

### Mechanisms of IOP elevation

*LMX1B* variants are known to disrupt drainage structure development and cause developmental and juvenile onset glaucoma (Lichter et al., 1997; Liu and Johnson, 2010; Pressman et al., 2000; Sawamura et al., 2014). These developmental changes lead to drainage structure abnormalities and IOP elevation. Our data clearly show that all *Lmx1b*^*V265D/+*^ eyes have structural abnormalities of their iridocorneal angle. B6 mutants had the greatest severity of angle abnormalities and the most severely dysregulated IOPs at younger ages. This suggests that developmental drainage structure abnormalities are important in IOP elevation in these mice. 129.*Lmx1b*^*V265D/+*^ mice have milder iridocorneal angle structural abnormalities with the vast majority of the angle being open, but they still develop elevated IOP. Mild iridocorneal angle defects are found in POAG patients with NPS, which is caused by *LMX1B* variants (Lichter et al., 1997; Vollrath et al., 1998). Thus, strain 129 mutants are a valuable resource to model IOP elevation in POAG due to *LMX1B* variants.

Although structural developmental changes cause early-onset elevated IOP in some mutants, IOP becomes high at older ages in other *Lmx1b* mutants. As mutant eyes have less functional drainage tissue to begin with, the remaining functional tissue may be more susceptible to damage with age, leading to later-onset IOP elevation. It is possible that mechanisms unrelated to structure or normal drainage-function are involved in *Lmx1b*-phenotypes during development or adult life (Gould et al., 2004). Mutants may have abnormal metabolism or suboptimal defense mechanisms against ongoing stressors (e.g. oxidative stress) leading to tissue demise and IOP elevation over time. In agreement with this, the majority of patients with *LMX1B* variants have primary open angle glaucoma (Sweeney et al., 2003). These patients develop IOP elevation at older ages and have an open drainage angle with no obvious structural abnormalities. The mechanisms by which *LMX1B* variants impact the function of drainage tissue in POAG are likely complex and require additional characterization.

In addition to IOP elevation, abnormally low IOP was found in *Lmx1b*^*V265D/+*^ mice on each strain background at various ages. Abnormally low IOP is observed in other mouse models with abnormal anterior segment development (Chang et al., 2001). One contributing factor could be dysgenesis of the ciliary body, which produces AqH (Chang et al., 2001). *Lmx1b* is expressed in the developing ciliary body (Pressman et al., 2000), and the *Lmx1b*^*V265D*^ allele could potentially cause dysfunction of AqH production. Additionally, the *Lmx1b*^*V265D*^ allele induces severe corneal phenotypes involving extensive stretching, ulceration, and perforation, that contribute to lower than normal IOP. B6.*Lmx1b*^*V265D/+*^ mice have the highest incidence of abnormally low IOP values and the most severely affected corneas, consistent with a role of corneal phenotypes in lowering IOP.

### Identifying the genetic modifiers

The genetic loci that modify glaucoma susceptibility in individuals with *LMX1B* variants are not known. The interactions between these loci are likely complex. We discovered a QTL on Chromosome 18 that predisposes *Lmx1b*^*V265D/+*^ mice to severe ocular abnormalities and IOP elevation. Future work is required to characterize specific modifiers, to understand disease risk of individuals with *LMX1B* mutations and provide molecular targets for therapies to treat IOP elevation and glaucoma. As the 69.5 megabase interval on Chr 18 contains approximately 442 protein-coding genes, we are currently unable to nominate specific candidates responsible for strain-specific differences in susceptibility. However, ongoing work is aimed at prioritizing positional candidates. To identify candidate modifier genes, we examined the Chr 18 QTL interval for regions that significantly differ between the B6 and strain 129 backgrounds. We discovered an interval around the Zinc finger E-box-binding homeobox 1 (*ZEB1*) locus that harbors several polymorphisms between B6 and 129 genomes, including predicted functional variants in *ZEB1* (Keane et al., 2011). Interestingly, *ZEB1* variants cause Fuch’s corneal endothelial dystrophy (FCED) (Gupta et al., 2015). In FCED, the corneal endothelial structure is disrupted causing corneal haze. Corneal haze differs significantly between B6 and strain 129 *Lmx1b* mutant mice at each examined age. Furthermore, *Zeb1* null mouse embryos have ocular developmental defects similar to *Lmx1b* mutants including iridocorneal adhesions (Liu et al., 2008). However, no links are yet established between *ZEB1* and IOP elevation or LMX1B. Therefore, although *ZEB1* is an intriguing candidate, the modifier interval requires further refinement before a specific locus can be identified. In conclusion, this study lays a strong foundation for better understanding mechanisms by which LMX1B contributes to glaucoma and for characterizing new therapeutic targets.

## Acknowledgements

The Authors would like to thank the Histology Services and Computational Services at The Jackson Laboratory, animal care staff at The Jackson laboratory and Columbia University, and Amy Bell for intraocular pressure measurements.

