## Supplemental figures and tables for "Genetic background modifies vulnerability to glaucoma related phenotypes in *Lmx1b* mutant mice"

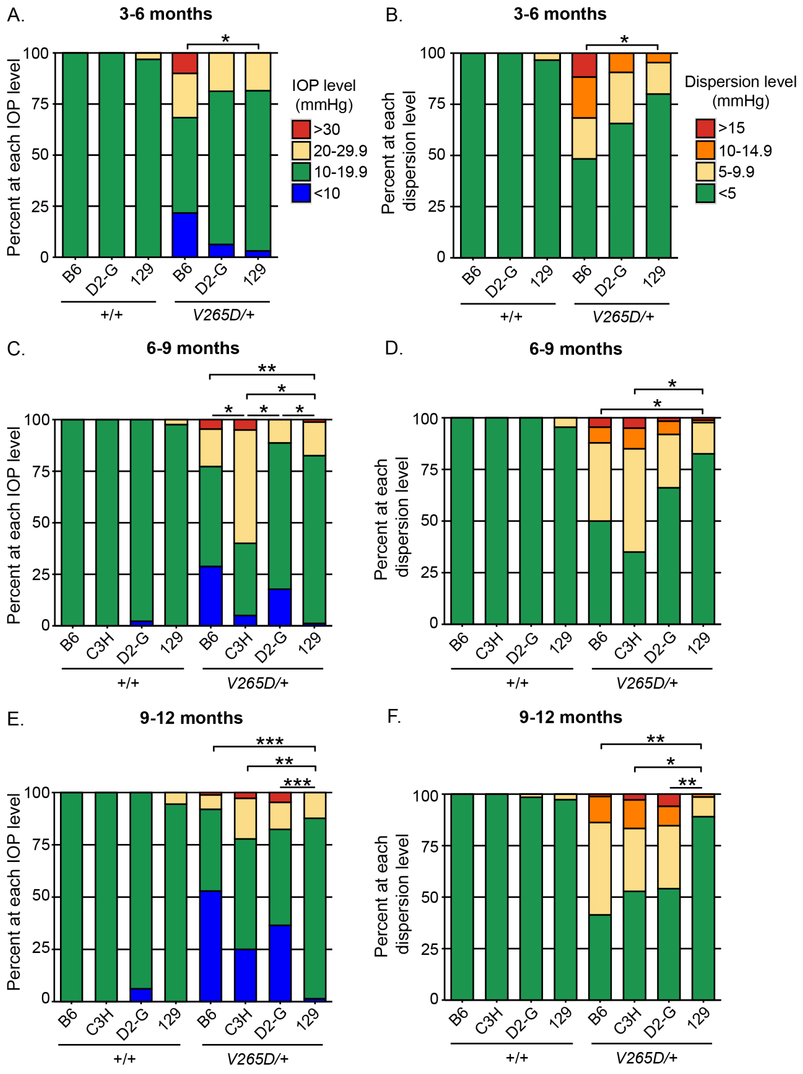


**Figure S1: Frequency distributions of binned IOP values.**

The data shown in Figure 4 were binned to help to visualize strain difference. **A,C,E** Binned IOP values. **B,D,F** Binned IOP dispersion values. * P<0.01, ** P<1E-05, *** P<1E-10 (see supplementary table 2 for exact P-values).

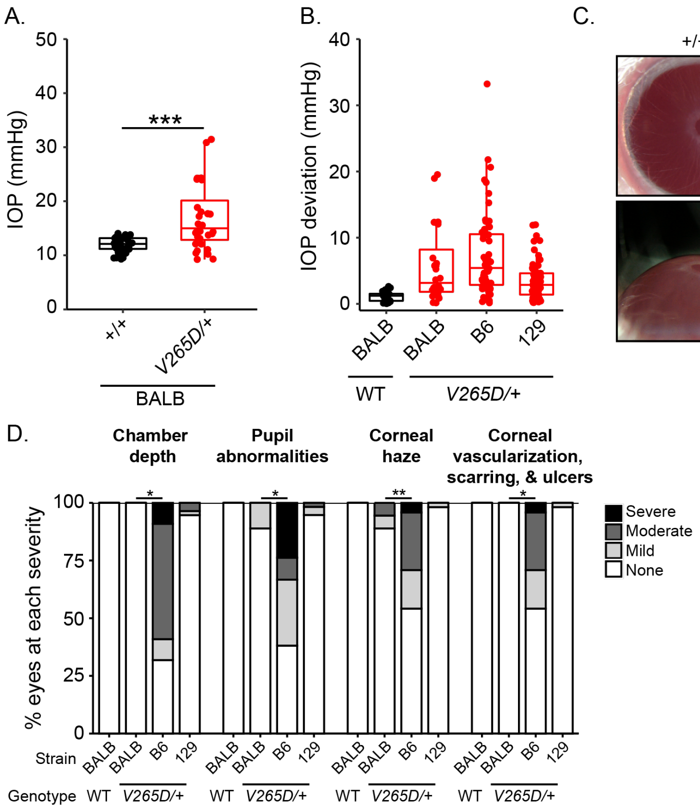


**Figure S2: The tyrosinase deficient BALB background does not exacerbate *Lmx1b* mutant phenotypes. (A and B)** Mutants on the BALB background have elevated IOP compared to WT controls ($p$ = 5.4E-05). BALB background does not increase the IOP deviation compared to other pigmented backgrounds (B6 and 129). **(C)** Representative slit-lamp photos of both front and side views from BALB WT and mutant eyes at 3 months of age. **(D)** BALB mutant mice rarely have anterior segment abnormalities and when present they were mild. See supplementary table 2 for P values. All mice were examined between 3 and 6 months of age. At least 16 eyes were examined in each group. * P<0.01; ** P<1.0E-05; *** P<1.0E-10 (see supplementary Table 1 for P values).

**Table S1:**

### A. Anterior chamber deepening *P* values

**3-5 months old**

| Group | B6 WT | B6 HET | D2-G HET | 129 HET | BALB HET |
|---|---|---|---|---|---|
| B6 WT | NA | | | | |
| B6 HET | 3.33E-07 | NA | | | |
| D2-G HET | NA | 0.0044 | NA | | |
| 129 HET | NA | 1.93E-08 | 2.76E-14 | NA | |
| BALB HET | NA | 1.92E-05 | 1.59E-07 | NS | NA |

**6-8 months old**

| Group | B6 WT | B6 HET | C3H HET | D2-G HET | 129 HET |
|---|---|---|---|---|---|
| B6 WT | NA | | | | |
| B6 HET | 2.32E-07 | NA | | | |
| C3H HET | NA | NS | NA | | |
| D2-G HET | NA | NS | NS | NA | |
| 129 HET | NA | 3.01E-08 | 0.00067 | 1.51E-10 | NA |

**10-12 months old**

| Group | B6 WT | B6 HET | C3H HET | D2-G HET | 129 HET |
|---|---|---|---|---|---|
| B6 WT | NA | | | | |
| B6 HET | 2.11E-16 | NA | | | |
| C3H HET | NA | NS | NA | | |
| D2-G HET | NA | NS | NS | NA | |
| 129 HET | NA | 1.08E-11 | 6.14E-18 | 2.26E-20 | NA |

### B. Pupil abnormalities *P* values

**3-5 months old**

| Group | B6 WT | B6 HET | D2-G HET | 129 HET | BALB HET |
|---|---|---|---|---|---|
| B6 WT | NA | | | | |
| B6 HET | 4.71E-05 | NA | | | |
| D2-G HET | NA | NS | NA | | |
| 129 HET | NA | 2.83E-07 | 3.10E-04 | NA | |
| BALB HET | NA | 4.98E-03 | NS | NS | NA |

**6-8 months old**

| Group | B6 WT | B6 HET | C3H HET | D2-G HET | 129 HET |
|---|---|---|---|---|---|
| B6 WT | NA | | | | |
| B6 HET | 4.55E-09 | NA | | | |
| C3H HET | NA | NS | NA | | |
| D2-G HET | NA | NS | NS | NA | |
| 129 HET | NA | 1.22E-10 | 4.41E-07 | 2.83E-07 | NA |

**10-12 months old**

| Group | B6 WT | B6 HET | C3H HET | D2-G HET | 129 HET |
|---|---|---|---|---|---|
| B6 WT | NA | | | | |
| B6 HET | 3.03E-18 | NA | | | |
| C3H HET | NA | NS | NA | | |
| D2-G HET | NA | NS | NS | NA | |
| 129 HET | NA | 5.83E-12 | 7.26E-15 | 8.23E-10 | NA |

### C. Corneal haze *P* values

**3-5 months old**

| Group | B6 WT | B6 HET | D2-G HET | 129 HET | BALB HET |
|---|---|---|---|---|---|
| B6 WT | NA | | | | |

| | | | | | |
|---|---|---|---|---|---|
| B6 HET | 6.20E-14 | NA | | | |
| D2-G HET | NA | 2.05E-10 | NA | | |
| 129 HET | NA | 6.78E-11 | NS | NA | |
| BALB HET | NA | 1.27E-09 | 7.61E-03 | NS | NA |

**6-8 months old**

| Group | B6 WT | B6 HET | C3H HET | D2-G HET | 129 HET |
|---|---|---|---|---|---|
| B6 WT | NA | | | | |
| B6 HET | 4.82E-15 | NA | | | |
| C3H HET | NA | 1.59E-05 | NA | | |
| D2-G HET | NA | 1.97E-08 | NS | NA | |
| 129 HET | NA | 2.42E-09 | NS | NS | NA |

**10-12 months old**

| Group | B6 WT | B6 HET | C3H HET | D2-G HET | 129 HET |
|---|---|---|---|---|---|
| B6 WT | NA | | | | |
| B6 HET | 2.69E-19 | NA | | | |
| C3H HET | NA | NS | NA | | |
| D2-G HET | NA | 3.80E-06 | 0.00121 | NA | |
| 129 HET | NA | 1.13E-06 | 0.00393 | NS | NA |

### D. Corneal vascularization, scarring, and ulcers *P* values

**3-5 months old**

| Group | B6 WT | B6 HET | D2-G HET | 129 HET | BALB HET |
|---|---|---|---|---|---|
| B6 WT | NA | | | | |
| B6 HET | 2.21E-04 | NA | | | |
| D2-G HET | NA | NS | NA | | |
| 129 HET | NA | 1.61E-06 | 2.96E-05 | NA | |
| BALB HET | NA | 3.49E-03 | 7.41E-03 | NS | NA |

**6-8 months old**

| Group | B6 WT | B6 HET | C3H HET | D2-G HET | 129 HET |
|---|---|---|---|---|---|
| B6 WT | NA | | | | |
| B6 HET | 4.05E-07 | NA | | | |
| C3H HET | NA | NS | NA | | |
| D2-G HET | NA | NS | NS | NA | |
| 129 HET | NA | 9.72E-08 | 4.19E-06 | 1.15E-04 | NA |

**10-12 months old**

| Group | B6 WT | B6 HET | C3H HET | D2-G HET | 129 HET |
|---|---|---|---|---|---|
| B6 WT | NA | | | | |
| B6 HET | 1.55E-12 | NA | | | |
| C3H HET | NA | NS | NA | | |
| D2-G HET | NA | NS | NS | NA | |
| 129 HET | NA | 7.25E-11 | 1.49E-15 | 3.71E-16 | NA |

NA, Not applicable (to test); NS, not significant

**Table S2:**

### A. IOP (Figure 4A-C)

**3-6 months old**

| Strain Background | *P* value (WT vs mutant) |
|---|---|
| B6 | NS |
| D2-G | 2.20E-04 |
| Strain 129 | 1.80E-05 |
| BALB | 5.44E-05 |

**6-9 months old**

| Strain Background | *P* value (WT vs mutant) |
|---|---|
| B6 | NS |
| C3H | 0.0032 |
| D2-G | 0.0097 |
| Strain 129 | 2.80E-04 |

**10-12 months old**

| Strain Background | *P* value (WT vs mutant) |
|---|---|
| B6 | 8.20E-07 |
| C3H | NS |
| D2-G | NS |
| Strain 129 | NS |

### B. IOP deviation (Figure 4D-F)

**3-6 months old *P* values**

| Group | B6 WT | D2-G WT | 129 WT | BALB WT | B6 HET | D2-G HET | 129 HET | BALB HET |
|---|---|---|---|---|---|---|---|---|
| B6 WT | NA | | | | | | | |
| D2-G WT | NS | NA | | | | | | |
| 129 WT | NS | NS | NA | | | | | |
| BALB WT | NS | NS | NS | NA | | | | |
| B6 HET | 3.50E-09 | NA | NA | NA | NA | | | |
| D2-G HET | NA | 3.10E-05 | NA | NA | 0.0068 | NA | | |
| 129 HET | NA | NA | 5.50E-06 | NA | 4.50E-05 | NS | NA | |
| BALB HET | NA | NA | NA | 4.16E-05 | NS | NS | NS | NA |

**6-9 months old *P* values**

| Group | B6 WT | C3H WT | D2-G WT | 129 WT | B6 HET | C3H HET | D2-G HET | 129 HET |
|---|---|---|---|---|---|---|---|---|
| B6 WT | NA | | | | | | | |
| C3H WT | NS | NA | | | | | | |
| D2-G WT | NS | NS | NA | | | | | |
| 129 WT | NS | NS | NS | NA | | | | |
| B6 HET | 6.90E-11 | NA | NA | NA | NA | | | |
| C3H HET | NA | 0.0003 | NA | NA | NS | NA | | |
| D2-G HET | NA | NA | 5.60E-08 | NA | NS | NS | NA | |
| 129 HET | NA | NA | NA | NS | 1.00E-05 | 0.009 | NS | NA |

**9-12 months old *P* values**

| Group | B6 WT | C3H WT | D2-G WT | 129 WT | B6 HET | C3H HET | D2-G HET | 129 HET |
|---|---|---|---|---|---|---|---|---|
| B6 WT | NA | | | | | | | |
| C3H WT | NS | NA | | | | | | |
| D2-G WT | NS | NS | NA | | | | | |
| 129 WT | NS | NS | NS | NA | | | | |
| B6 HET | 1.80E-17 | NA | NA | NA | NA | | | |
| C3H HET | NA | 8.8594E-07 | NA | NA | NS | NA | | |
| D2-G HET | NA | NA | 7.4831E-12 | NA | NS | NS | NA | |
| 129 HET | NA | NA | NA | NS | 3.0677E-11 | 2.4517E-05 | 5.6186E-08 | NA |

### C. IOP distribution (Figure S1A,C,E)

**3-6 months old *P* values**

| Group | B6 WT | D2-G WT | 129 WT | B6 HET | D2-G HET | 129 HET |
|---|---|---|---|---|---|---|
| B6 WT | NA | | | | | |
| D2-G WT | NS | NA | | | | |
| 129 WT | NS | NS | NA | | | |
| B6 HET | 9.00E-12 | NA | NA | NA | | |

| | | | | | | |
|---|---|---|---|---|---|---|
| D2-G HET | NA | 6.60E-04 | NA | NS | NA | |
| 129 HET | NA | NA | NS | 3.50E-05 | NS | NA |

**6-9 months old *P* values**

| Group | B6 WT | C3H WT | D2-G WT | 129 WT | B6 HET | C3H HET | D2-G HET | 129 HET |
|---|---|---|---|---|---|---|---|---|
| B6 WT | NA | | | | | | | |
| C3H WT | NS | NA | | | | | | |
| D2-G WT | NS | NS | NA | | | | | |
| 129 WT | NS | NS | NS | NA | | | | |
| B6 HET | 1.50E-12 | NA | NA | NA | NA | | | |
| C3H HET | NA | 4.00E-04 | NA | NA | NS | NA | | |
| D2-G HET | NA | NA | 6.60E-04 | NA | NS | NS | NA | |
| 129 HET | NA | NA | NA | NS | 2.10E-07 | 1.10E-04 | 0.0012 | NA |

**9-12 months old *P* values**

| Group | B6 WT | C3H WT | D2-G WT | 129 WT | B6 HET | C3H HET | D2-G HET | 129 HET |
|---|---|---|---|---|---|---|---|---|
| B6 WT | NA | | | | | | | |
| C3H WT | NS | NA | | | | | | |
| D2-G WT | NS | NS | NA | | | | | |
| 129 WT | NS | NS | NS | NA | | | | |
| B6 HET | 2.20E-16 | NA | NA | NA | NA | | | |
| C3H HET | NA | 0.000106 | NA | NA | NS | NA | | |
| D2-G HET | NA | NA | 7.49E-10 | NA | NS | NS | NA | |
| 129 HET | NA | NA | NA | NS | 1.80E-14 | 3.46E-05 | 6.65E-10 | NA |

### D. IOP deviation distribution (Figure S1B,D,F)

**3-6 months old *P* values**

| Group | B6 WT | D2-G WT | 129 WT | B6 HET | D2-G HET | 129 HET |
|---|---|---|---|---|---|---|
| B6 WT | NA | | | | | |
| D2-G WT | NS | NA | | | | |
| 129 WT | NS | NS | NA | | | |
| B6 HET | 2.81E-11 | NA | NA | NA | | |
| D2-G HET | NA | 3.57E-04 | NA | NS | NA | |
| 129 HET | NA | NA | NS | 1.38E-04 | NS | NA |

**6-9 months old *P* values**

| Group | B6 WT | C3H WT | D2-G WT | 129 WT | B6 HET | C3H HET | D2-G HET | 129 HET |
|---|---|---|---|---|---|---|---|---|
| B6 WT | NA | | | | | | | |
| C3H WT | NS | NA | | | | | | |
| D2-G WT | NS | NS | NA | | | | | |
| 129 WT | NS | NS | NS | NA | | | | |
| B6 HET | 5.56E-12 | NA | NA | NA | NA | | | |
| C3H HET | NA | 3.97E-04 | NA | NA | NS | NA | | |
| D2-G HET | NA | NA | 1.13E-05 | NA | NS | NS | NA | |
| 129 HET | NA | NA | NA | NS | 8.25E-05 | 5.08E-05 | NS | NA |

**9-12 months old *P* values**

| Group | B6 WT | C3H WT | D2-G WT | 129 WT | B6 HET | C3H HET | D2-G HET | 129 HET |
|---|---|---|---|---|---|---|---|---|
| B6 WT | NA | | | | | | | |
| C3H WT | NS | NA | | | | | | |
| D2-G WT | NS | NS | NA | | | | | |
| 129 WT | NS | NS | NS | NA | | | | |
| B6 HET | 1.49E-15 | NA | NA | NA | NA | | | |
| C3H HET | NA | 9.41E-05 | NA | NA | NS | NA | | |

| | | | | | | | | |
|---|---|---|---|---|---|---|---|---|
| D2-G HET | NA | NA | 5.71E-10 | NA | NS | NS | NA | |
| 129 HET | NA | NA | NA | NS | 1.06E-09 | 9.47E-05 | 6.20E-06 | NA |

NA, Not applicable (to test); NS, not significant
